# Advancing age grading techniques for *Glossina morsitans morsitans*, vectors of African trypanosomiasis, through mid-infrared spectroscopy and machine learning

**DOI:** 10.1101/2024.02.19.579508

**Authors:** Mauro Pazmiño-Betancourth, Ivan Casas Gómez-Uribarri, Karina Mondragon-Shem, Simon A Babayan, Francesco Baldini, Lee Rafuse Haines

**Author notes:** These authors equally supervised the work (Francesco Baldini, Lee Rafuse Haines).

## Abstract

Tsetse are the insects responsible for transmitting African trypanosomes, which cause sleeping sickness in humans and animal trypanosomiasis in wildlife and livestock. Knowing the age of these flies is important when assessing the effectiveness of vector control programs and modelling disease risk. However, current methods to assess fly age are labour-intensive, slow, and often inaccurate as skilled personnel are in short supply. Mid-infrared spectroscopy (MIRS), a fast and cost-effective tool to accurately estimate several biological traits of insects, offers a promising alternative. This is achieved by characterising the biochemical composition of the insect cuticle using infrared light coupled with machine learning algorithms to estimate the traits of interest.

We tested the performance of MIRS in estimating tsetse sex and age for the first time using spectra obtained from their cuticle. We used 541 insectary-reared *Glossina m. morsitans* of two different age groups for males (5 and 7 weeks) and three age groups for females (3 days, 5 weeks, and 7 weeks). Spectra were collected from the head, thorax, and abdomen of each sample. Machine learning models differentiated between male and female flies with a 96% accuracy and predicted the age group with 94% and 87% accuracy for males and females, respectively. The key infrared regions important for discriminating sex and age classification were characteristic of lipid and protein content. Our results support the use of MIRS as a fast and accurate way to identify tsetse sex and age with minimal pre-processing. Further validation using wild-caught tsetse can pave the way for this technique to be implemented as a routine surveillance tool in vector control programmes.

**Author summary:** Male and female tsetse transmit the parasites that cause sleeping sickness in humans and nagana in livestock. To control these diseases, knowing the age of these flies is important, as it helps evaluate the efficacy of control measures and assess disease risk. However, current age-grading methods are laborious, often unreliable, and in the case of male tsetse, highly inaccurate. This study explores a novel approach that uses mid-infrared spectroscopy (MIRS) to estimate the age of individual tsetse. Machine learning can detect signatures in MIRS that help identify the composition of a fly’s cuticle, which differs between sexes and changes as they age.

We trained machine learning models that distinguished male from female flies with 96% accuracy and predicted the correct age group with 94% accuracy for males and 87% accuracy for females. MIRS offers a fast and reliable way to identify tsetse sex and age with minimal preparation. If this method is successfully validated with wild flies, it holds the potential to vastly increase the accuracy of the way we monitor and combat these disease-carrying insects, thus offering significant advantages in our efforts to control them.

## Introduction

Tsetse are blood-feeding flies that can transmit trypanosome parasites of human and animal concern. There are two parasite species that cause Human African Trypanosomiasis (HAT), or sleeping sickness, and infected patients can die if they do not receive treatment. The promising decline of cases in endemic areas[1] in recent years is due to ongoing disease and vector control efforts, but continued support is critical to ensure the success of disease elimination programmes. However, Rhodesiense HAT (the more severe form) is still a concern due to livestock and wildlife forming part of its transmission cycle. Animal African trypanosomiasis (AAT) affects wildlife and domestic animals, causing three million cattle deaths/year with agricultural losses nearing US$ 5 billion/year[2]. Both female and male tsetse can transmit trypanosomes, but only adult flies older than 20 days post-emergence that have ingested blood from a parasite-infected host can be infectious. Tsetse age is therefore crucial for estimating transmission risk and the efficacy of vector control programmes. Accurate age grading in the field is crucial for disease monitoring and evaluation operations. An effective vector control intervention, which does not discriminate against age, overall will reduce the average age of tsetse populations. For example, if in an area of ongoing vector control only young flies are caught, this suggests newly emerged flies in the area, whereas capturing older flies either indicates fly reinvasion from outside the intervention zone or intervention failure.

Tsetse age grading for female flies currently relies on performing a labour-intensive ovarian dissection, which requires the use of a microscope and an experienced dissector. Female tsetse give birth to a larva every 9 days[3] throughout life, and the four ovarioles develop in a specific, predictable sequence; as each egg descends into the uterus, it leaves behind a scar (named ‘relic’) that can be microscopically identified[4]. No new relics are created after the 4th ovarian cycle, thus limiting the confidence of this method in flies older than seven weeks[4]. Furthermore, factors such as nutritional stress[5], tsetse strain[6] and temperature[7] can affect the length of this 9-day process, and even with adjustments, the method can be imprecise. Ovarian dissections are time consuming and need to be performed while the tsetse is still ‘fresh’, and tissues maintain their form. After death, flies quickly become dehydrated and age grading is no longer possible by this method. This makes it difficult to process large numbers of flies when monitoring control interventions.

The current situation is worse for male tsetse, as there are no dependable methods for age-grading them. Wing fray analysis in either wild male[8] or female flies is unreliable as artifacts can be introduced through trapping protocols. Other approaches like tsetse eye pigment (pteridine) analysis[9] and gene expression [10] are too complex or costly for routine use in field settings. Thus, all current age-grading methods are either too imprecise, laborious, or expensive.

Mid-infrared spectroscopy (MIRS) has proven to be a versatile technique for determining mosquito age and species in both insectary-reared and field-collected mosquitoes[11–14]. MIRS quantifies the energy a molecule absorbs based on its molecular vibrations[15,16]. As the insect surface is covered with a complex mixture of cuticular proteins, polysaccharides, wax and other lipids, this tool provides a way to detect the differences between different samples. The chemical composition of male and female cuticles, as well as different species-specific signatures, can be resolved alongside more transient aspects such as cuticular changes over time[17]. Scanning a dried insect sample with MIRS is fast (1-2 minutes), and when combined with the use of machine learning (ML) algorithms, it provides a powerful toolbox for researchers to rapidly assess vector populations with minimum sample processing and high accuracy.

In this study, we use ML to estimate the age and sex from MIRS of different fly tissues collected from insectary-reared tsetse (*Glossina morsitans morsitans)* of known age and sex. We also identified the regions of the tsetse mid-infrared spectrum associated with age and sex, to elucidate the biological basis of our model predictions.

## Methods

### Tsetse rearing

An age-stratified colony of *Glossina morsitans morsitans* Westwood, established in 2004 at the Liverpool School of Tropical Medicine (LSTM), UK, was daily maintained under the following conditions: 26 – 28 °C, 68 – 78 % humidity and a 12 h/12 h light/dark cycle. Tsetse were fed three times a week on sterile defibrinated horse blood (TCS Biosciences Ltd, Buckingham, UK) using a silicon membrane feeding system.

### Tsetse sampling strategy and desiccation

Young, unmated female flies were first collected from emerging pupal pots as male emergence is delayed, and male collection was timed after the females had emerged. Both teneral (unfed, newly emerged) female and male collections were isolated from each other to prevent potential cuticular contamination with contact sex pheromones (cuticular hydrocarbon) during mating[18].

We collected 354 female and 187 male teneral tsetse from the LSTM colony in total for analysis. At specific ages, tsetse were killed with chloroform-soaked cotton, placed on a thin layer of cotton wool inside a 15 ml falcon tube half-filled with silica gel beads, sealed and then stored at 4°C until required. Desiccated tsetse were transferred to 96-well plates in preparation for shipping to the University of Glasgow. Upon analysis, dried flies were dissected into three sections: head, thorax and abdomen using dissection tweezers.

### Infrared Spectroscopy

Spectra from individual heads, thoraces and abdomens were taken by Attenuated Total Reflection (ATR) FT-IR spectroscopy using a Bruker ALPHA II spectrometer equipped with a Globar lamp, a deuterated L-alanine doped triglycene sulphate (DLaTGS) detector, a Potassium Bromide (KBr) beam splitter, and a diamond ATR accessory (Bruker Platinum ATR Unit A225). Twenty-four scans were collected at room temperature between 4000 and 400 cm^−1^ with 4 cm^−1^ resolution per sample. When measuring the tsetse samples, we made efforts to avoid practices that introduce sources of bias such as: not always measuring first young and then old samples, or first females and then males. Low-quality spectra were discarded using a custom script designed for mosquito spectra [11,19].

### Machine learning analysis

Spectra were centred around a mean of 0 and scaled to a standard deviation of 1 prior to any analysis. Uniform Manifold Approximation and Projection (UMAP) was applied for clustering analysis. Sex and age groups were binarized using one hot encoding[20]. First, we shuffled and split the dataset into the training (80%) and test sets (20%), stratified by sex and age groups (Supplementary Material Table S1). The training set was used to compute baseline performance of four machine learning algorithms: Logistic Regression (LR), Random Forest (RF), Support vector machine (SVC) and Classification and Regression Tree (CART) using 10-fold cross validation and default parameter settings on the training set. Additionally, a permutation score test was performed to evaluate if there was a dependency between the features (absorbance of each wavenumber) and classes (sex and age groups) (Supplementary Material Fig S1). The best model was then optimized using hyperparameter tuning, which consists in choosing a set of optimal values for the model hyperparameters to maximize its performance. The remaining 20% of the data (the test set) was used for the final evaluation of the optimized models. The individual metrics used to evaluate the models were accuracy, sensitivity, and specificity. Machine learning was performed using Python 3.10 and scikit-learn 1.2.2.

### Data and code availability

The infrared spectral data generated for this study have been deposited in the Enlighten database and are available at http://dx.doi.org/10.5525/gla.researchdata.1564.

All code to reproduce the machine learning analysis and figures is available at https://github.com/maurocolapso/Pazmino_TsetseMIRS_2023.git

## Results

### Optimization of tsetse desiccation

To understand how long it took for tsetse in different nutritional states to fully dehydrate (which is key to avoid the noise in the spectra caused by the water signal), we placed individual tsetse into 15ml tubes containing a deep layer of silica gel under a thin cap of cotton wool. Fly weight loss was daily recorded until it stabilised. Unfed flies rapidly desiccated within 24h, while fully engorged, bloodfed male and female flies took over three days to dehydrate the water-rich meal. Based on this data we adopted a standardized ∼72h of desiccation on silica for all flies subjected to MIRS analysis (**Error! Reference source not found.**)

### Differences between tsetse tissues

Initial tests focused on finding the best body regions or tissues to give a high signal clarity when doing spectrometric readings, as the large size tsetse presented novel logistical challenges. Because wild-caught flies are likely to acquire foreign hydrocarbons from mating, blood feeding, or the resting environment, we sampled zones of the cuticle expected to show the least contamination (**Error! Reference source not found.**).

We further investigated the variation between spectra of different tissues. Spectra from fly abdomens differed substantially those from heads and thoraces (Fig 2A), showing lower intensity and a higher variability, especially in the 1800 to 900 cm^-1^ region (Fig 2B). Moreover, visual inspection of the abdomens indicated that despite ∼60 days in a sealed anhydrous environment, complete desiccation was not achieved, particularly if the fly had ingested a large blood volume prior to collection. This residual horse blood and water could be driving the greater variability of the abdominal spectra compared to the other tissues. On top of that, previous work in other insects showed the thorax as a target tissue for MIRS. Consequently, we decided to focus our analysis on the spectra obtained from heads and thoraces only. A total of 1071 spectra were obtained by scanning the heads and lateral part of the thoraces of 541 flies of different ages (Fig 1, Table *2*).

**Fig 1.**
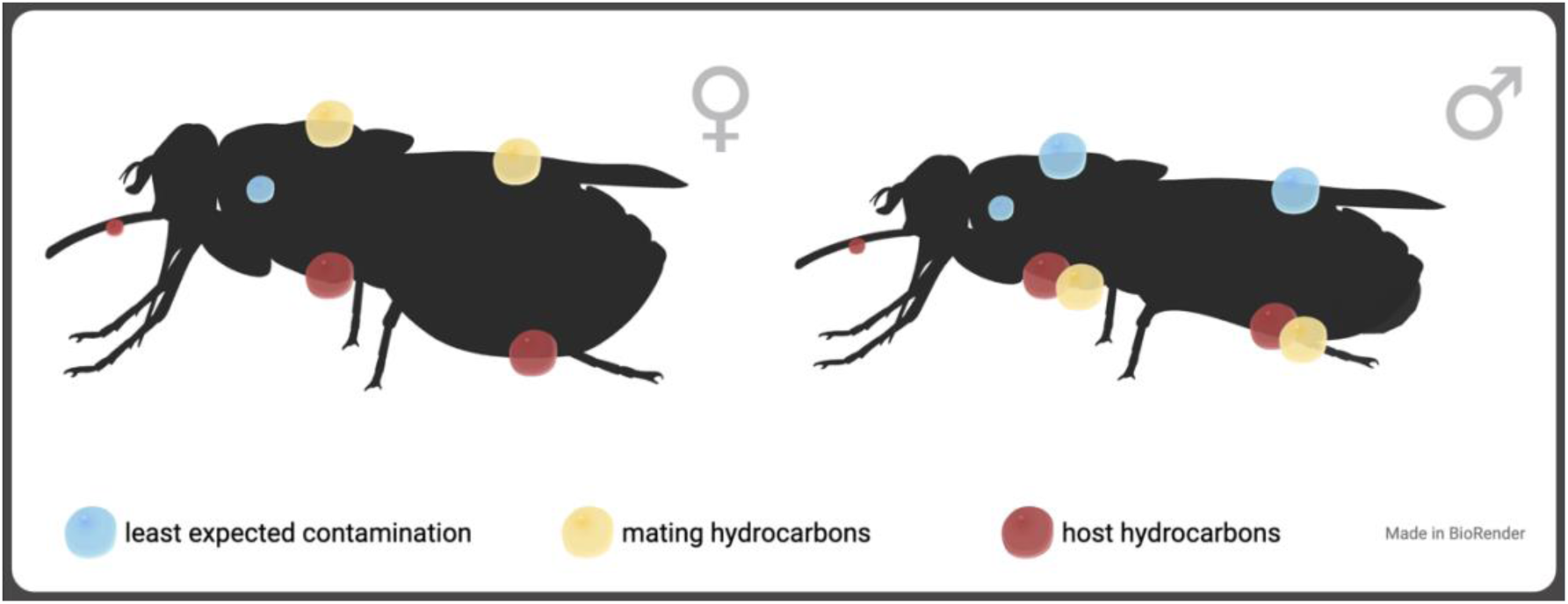
Tsetse biology and ecology suggest the heads and dorsal side of male tsetse or the lateral side of the thorax in both sexes would be the best areas (blue circles) to detect individual cuticular hydrocarbons.

**Fig 2.**
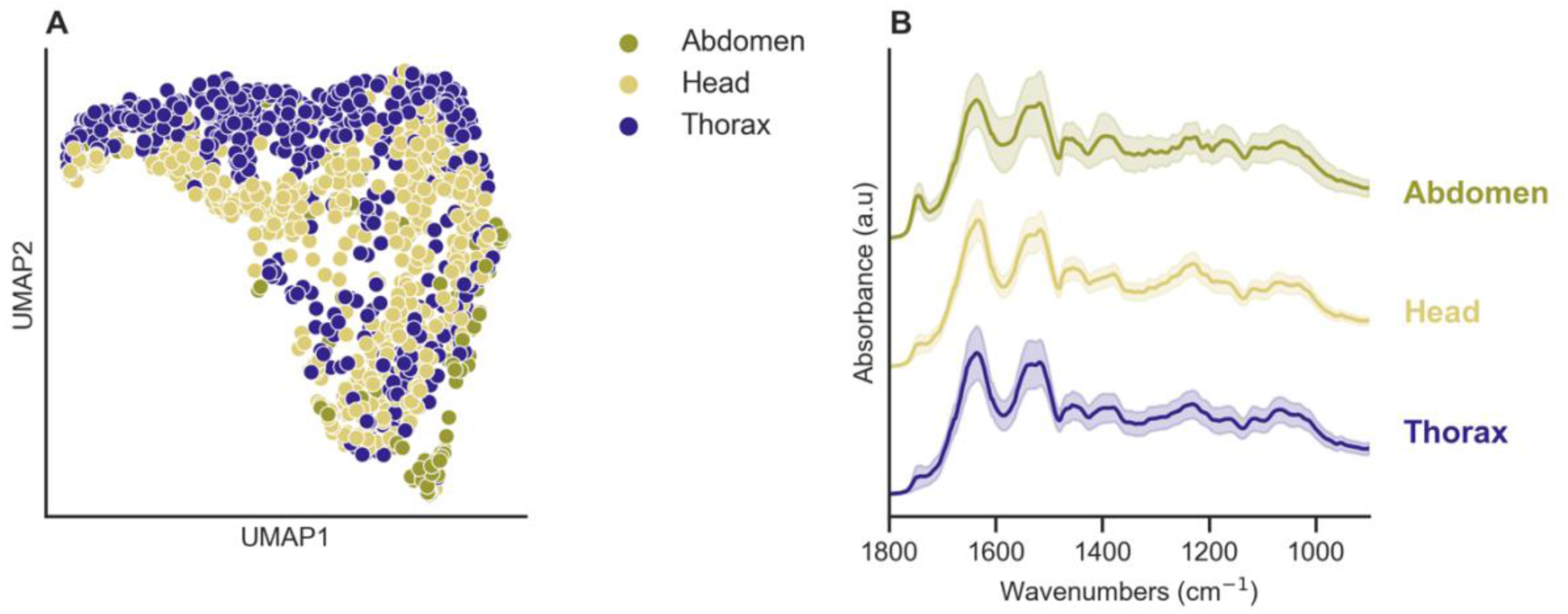
Spectra comparison from the abdomen, head, and thorax. A) Uniform Manifold Approximation and Projection (UMAP) of the abdomens, heads, and thoraces showed that the spectra collected from abdomens formed a separate cluster. Abdomens (olive green), head (yellow), thorax (purple) **B)** High variability of the spectra from abdomens (olive green line) showed that the sources of those inconsistencies were of low intensity and great variability at some wavelengths (primarily in the 1800 to 900 cm^-1^ region) compared to spectra from head (yellow line) and thorax (purple line). Spectra shown in panel B have been manually shifted across the Y-axis for ease of comparison.

**Table 1.**
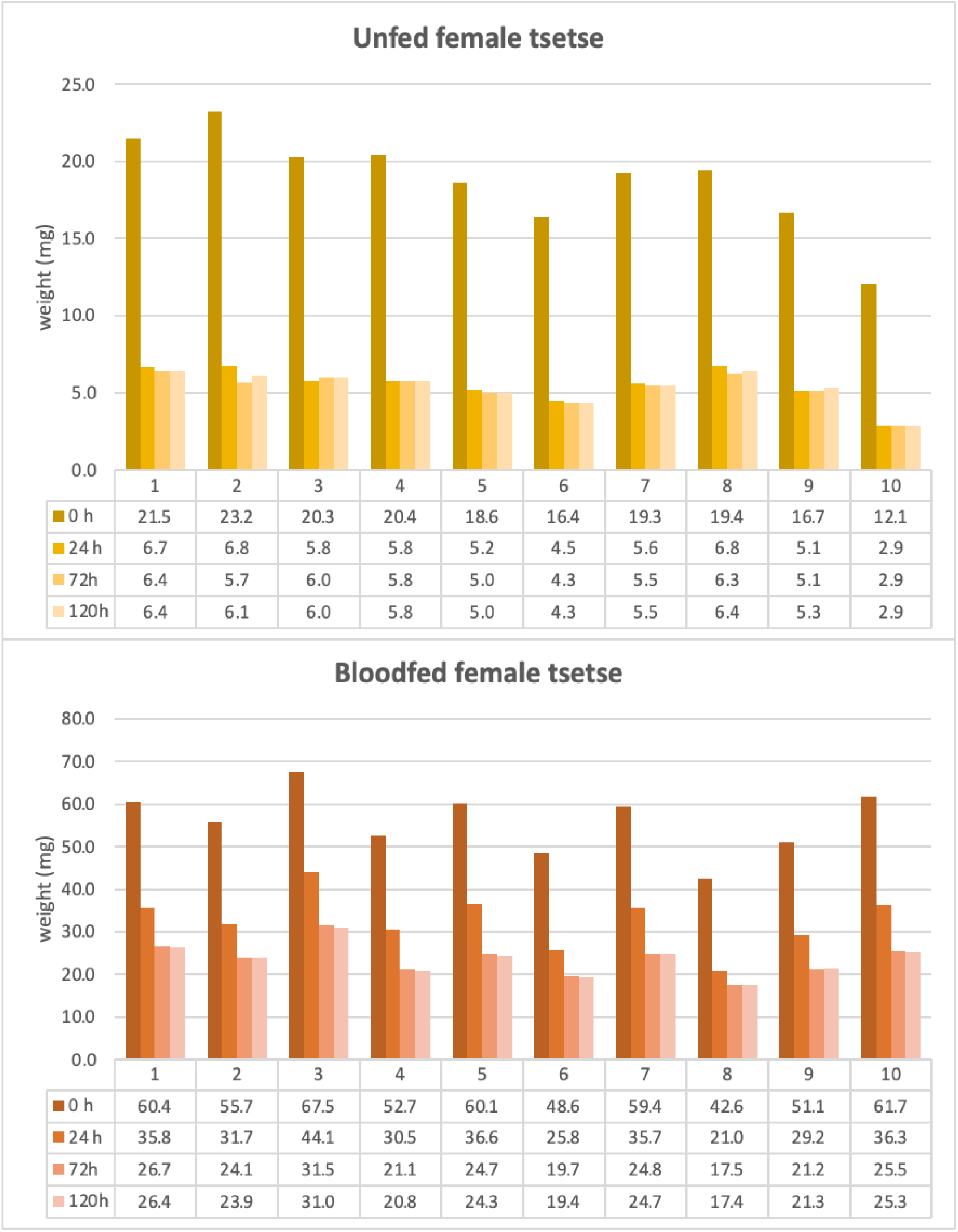
Desiccation time test for unfed and bloodfed female and male tsetse.

**Table 2.** Summary of aggregated samples sizes.

| Sex | Age | Tissue | # spectra | # samples |
| --- | --- | --- | --- | --- |
| Female | 3 days | Head | 133 | 136 |
|  |  | Thorax | 136 |  |
|  | 5 weeks | Head | 92 | 96 |
|  |  | Thorax | 96 |  |
|  | 7 weeks | Head | 120 | 122 |
|  |  | Thorax | 122 |  |
| Male | 5 weeks | Head | 94 | 94 |
|  |  | Thorax | 93 |  |
|  | 7 weeks | Head | 93 | 93 |
|  |  | Thorax | 92 |  |
| Total number of samples |  |  | 1071 | 541 |

### Differences between sexes and age groups

We used the unsupervised machine learning algorithm Uniform Manifold Approximation and Projection (UMAP) to investigate whether the spectra from fly heads (Fig 3 A-C) and thoraces (Fig 3 D-F) differed between flies of different sex and age. Most of the male flies produced different spectra than females, with the thorax showing clearer clusters with fewer samples overlapping between them (**Fig 3 A** and D). For age groups, there were not clusters in males regardless the tissue (**Fig 3 B** and E). In females, there were a distinct cluster composed of old flies (5 and 7 weeks) when using the thorax, however, there was a high overlap between samples from different age groups (**Fig 3 C** and F). These results show that MIRS contains biochemical information associated with sex and age as expected from relative changes in the cuticular composition of tsetse.

**Fig 3.**
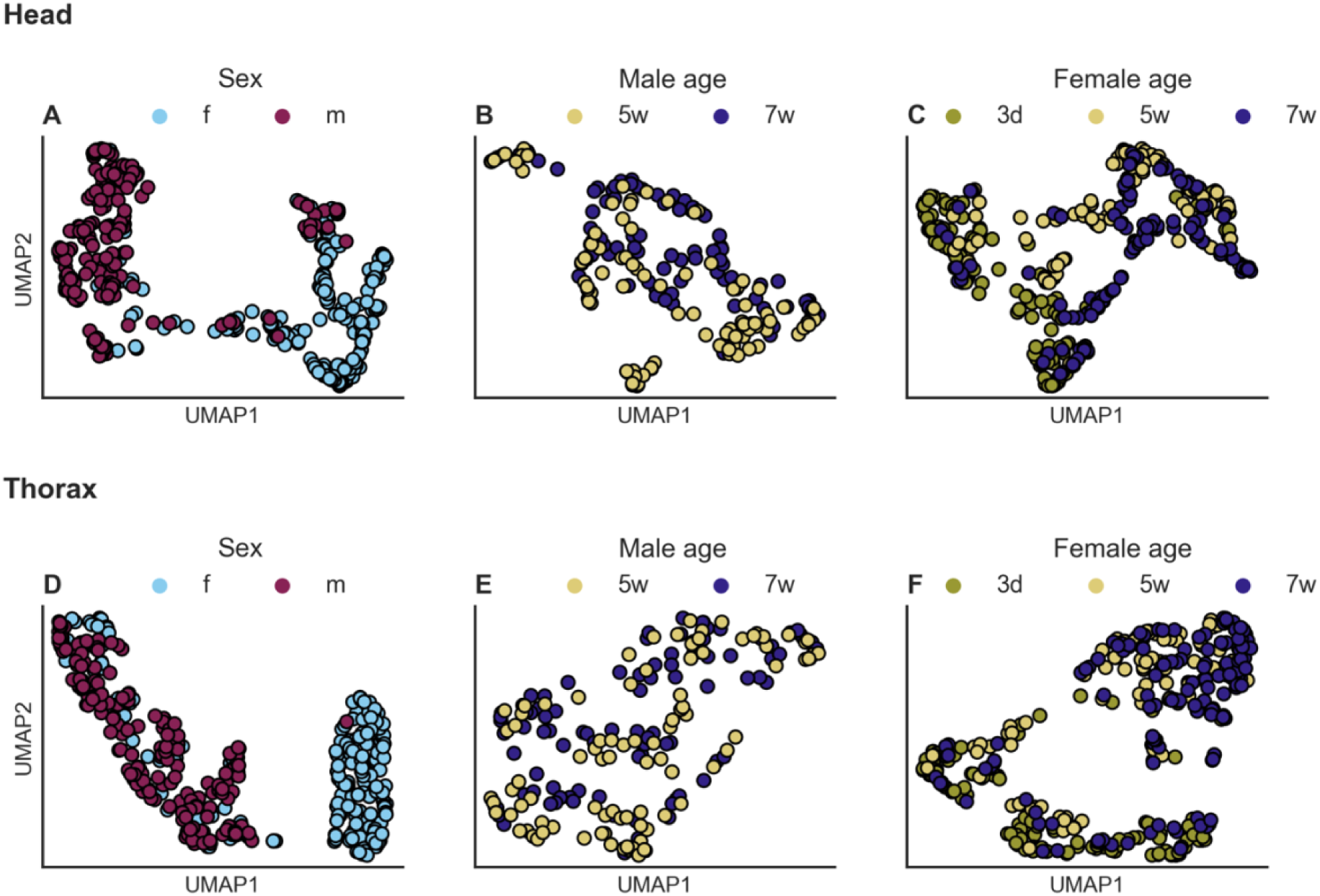
MIRS spectra according to tsetse sex and age from specific tissues. Unsupervised clustering of MIRS measurements using Uniform Manifold Approximation and Projection of MIRS in two-dimensional space using the heads and thorax. Samples are coloured by: **A, D**) sex (females: blue, males: purple). **B, E**) Males coloured by age (5 weeks: yellow, 7 weeks: dark blue). **C, F**) Females coloured by age (3 days: olive green, 5 weeks: yellow, 7 weeks: dark blue)

### Sex and age prediction using the complete spectral data

To identify tsetse sex and age-specific patterns within our MIRS dataset, we compared logistic regression (LR), Random Forest (RF), support vector machine (SVC) and the Classification and Regression Tree (CART) algorithms. Among these, LR had the highest accuracy (Fig 4A) when estimating the sex of five- and seven-week-old flies. Training accuracy was 94% when using both head (Fig 4A) and thorax (Supplementary Material Table S2). Similar accuracies were obtained in the test set (head = 99%, thorax = 94%, Supplementary Material Table S2). Logistic Regression was also the most accurate algorithm for identifying age groups among flies of the same sex (Fig 4C). For males, the thorax was marginally better at age prediction with an accuracy of 88% compared to 85% for the head (Supplementary Material Table S2). Similar performance was found in the test set with 92% and 89% for thorax and head, respectively (Supplementary Material Table S2). In females, even though there was some difference in accuracy between the head and thorax on the training set (head = 86%, thorax = 92%), accuracy on the test set was similar for both tissues at 93% (Supplementary Material Table S2). While these initial results suggest that infrared spectra could be used to predict key biological traits of tsetse flies, further analysis of model coefficients suggested that the predictions were being based mostly on flat regions of the spectra, between 4000 – 3750 cm^-1^ and 2250 – 1800 cm^-1^ (Fig 4 B, D and E), which are unlikely to contain biochemical information associated with insect cuticle[11] and are primarily used to monitor the presence of CO_2_ in the environment[15]. This phenomenon was observed with all predictive algorithms regardless of what tissue was used. To further investigate this, we applied the framework by Eid et al. [21]. Briefly, we divided the spectrum into three parts: two regions known to contain vibrations from key chemical bonds (3500 – 2500 cm^-1^ and 1800 – 600 cm^-1^) and one region where no chemical information associated with insect cuticle is expected (2500 – 1800 cm^-1^). We then compared the accuracy of four algorithms: Logistic regression, SVM with two kernels (RBF and linear) and Random Forest on each region. While the biochemical fingerprint regions (3500 – 2500 cm^-1^, 1800 – 600 cm^-1^) gave variable prediction accuracies (60 – 96%), when using the region with no chemical information associated with insect cuticle (2500 – 1800 cm^-1^), two algorithms (Logistic Regression and SVM with a linear kernel) could still predict different traits with high accuracy (83 – 94%), indicating possible overfitting (Supplementary Table S3). To produce more generalisable models, we therefore chose to base our predictions on the spectral region of 1800 – 600 cm^-1^, which is known to contain the most relevant biochemical information in insects[12–14].

**Fig 4.**
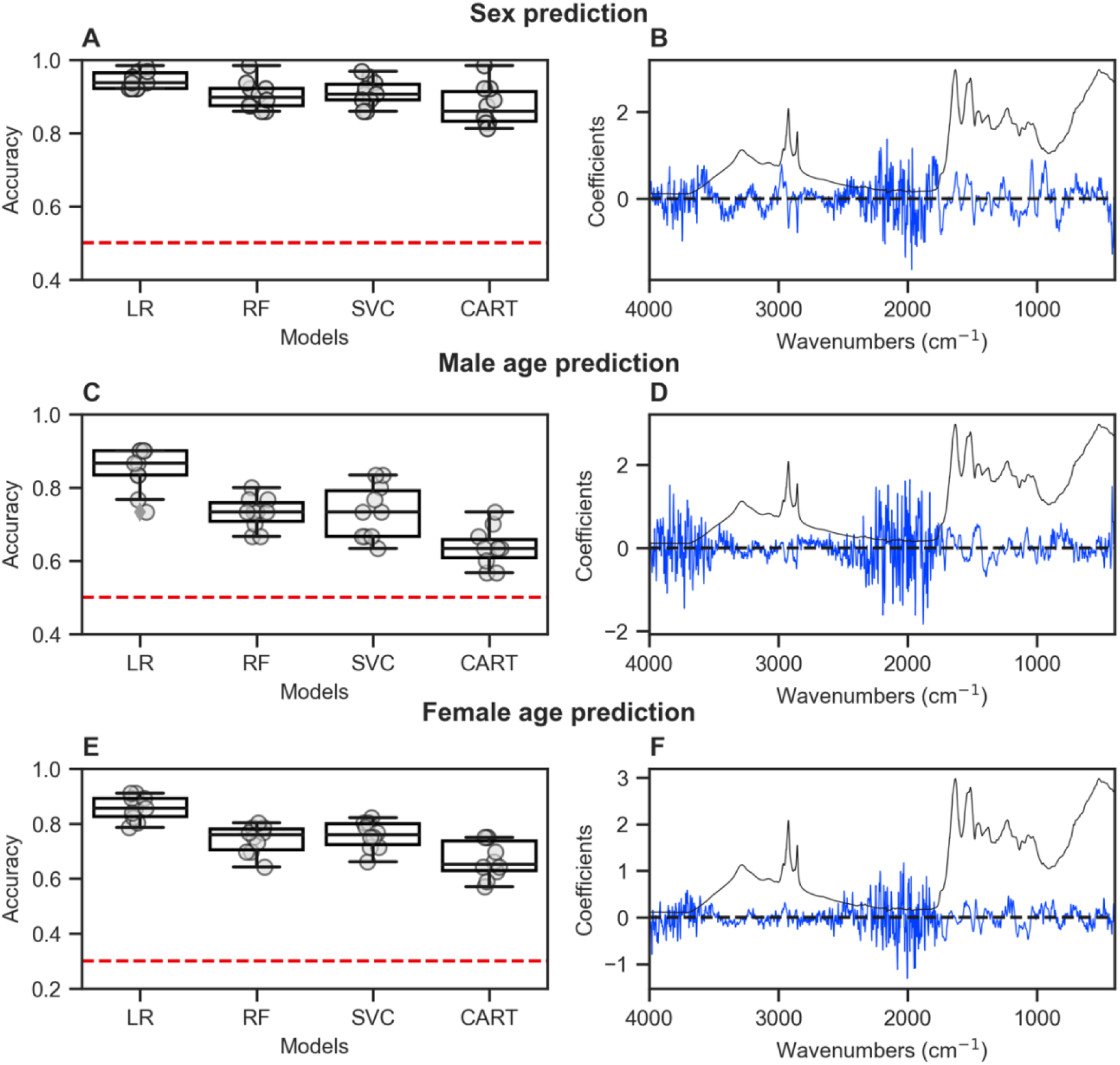
Prediction of tsetse sex and age using MIRS. Model performance on the training set of various ML models (LR: Logistic regression, RF: random forest, SVC: support vector machine and CART: decision tree classifier) for sex and age prediction using the heads of tsetse (**A, C, E**). Boxplots show the distribution of accuracies using 10-fold cross-validation. The horizontal dashed red line indicates a 0.5 accuracy for binary predictions (**A, C**) and 0.3 for a three-class prediction (**E**). Coefficients of the best model (blue line) plotted against the mean spectra of tsetse (**B, D, F**) show how the model relies on the 4000 – 3500 and 2500 –1800 cm^-1^ regions for prediction, which are lacking key biological information.

### Sex and age prediction using the biochemical fingerprint region of the spectra

When considering only the spectral region from 1750 - 600 cm^-1^, the accuracy of predicting fly sex and age marginally declined regardless of the algorithm used for analysis (Supplementary Material Fig S2). Logistic regression was able to clearly predict sex using the spectra from the heads with an accuracy of 96% (Fig 5A). The most informative wavenumbers appeared around the 1000 and 800 cm^-1^ areas of the spectra (Fig 6). Logistic regression was able to predict 5-week vs. 7-week-old males using spectral data from the head with an accuracy of 95% (Fig 5B). Most of the coefficients used by this model were in the 1636 and 1400 cm^-1^ region (Fig 6 Fig 6). Finally, age prediction in females was better when using the thorax. Young teneral flies were identified by the model with over 90% accuracy and older flies with 80% accuracy (Fig 5F). Like males, the important wavenumbers were in the same range, the 1750 to 1450 cm^-1^ 800 to 600 cm^-1^region (Fig 6). However, when using spectral data from the head, the model struggled to identify the 5-week-old age group; an accuracy of 58% (Fig 5C) was obtained, which was likely influenced by several 5-week-old samples being misclassified as 3-day-olds. A summary of the performance of Logistic Regression is shown in Table 3 and wavenumber importance and their assignments are presented in Supplementary Table S4. These results suggest that MIRS-ML is a promising approach when using the tsetse head or thorax to reliably produce quality spectra for sex and age prediction of laboratory-reared flies.

**Fig 5.**
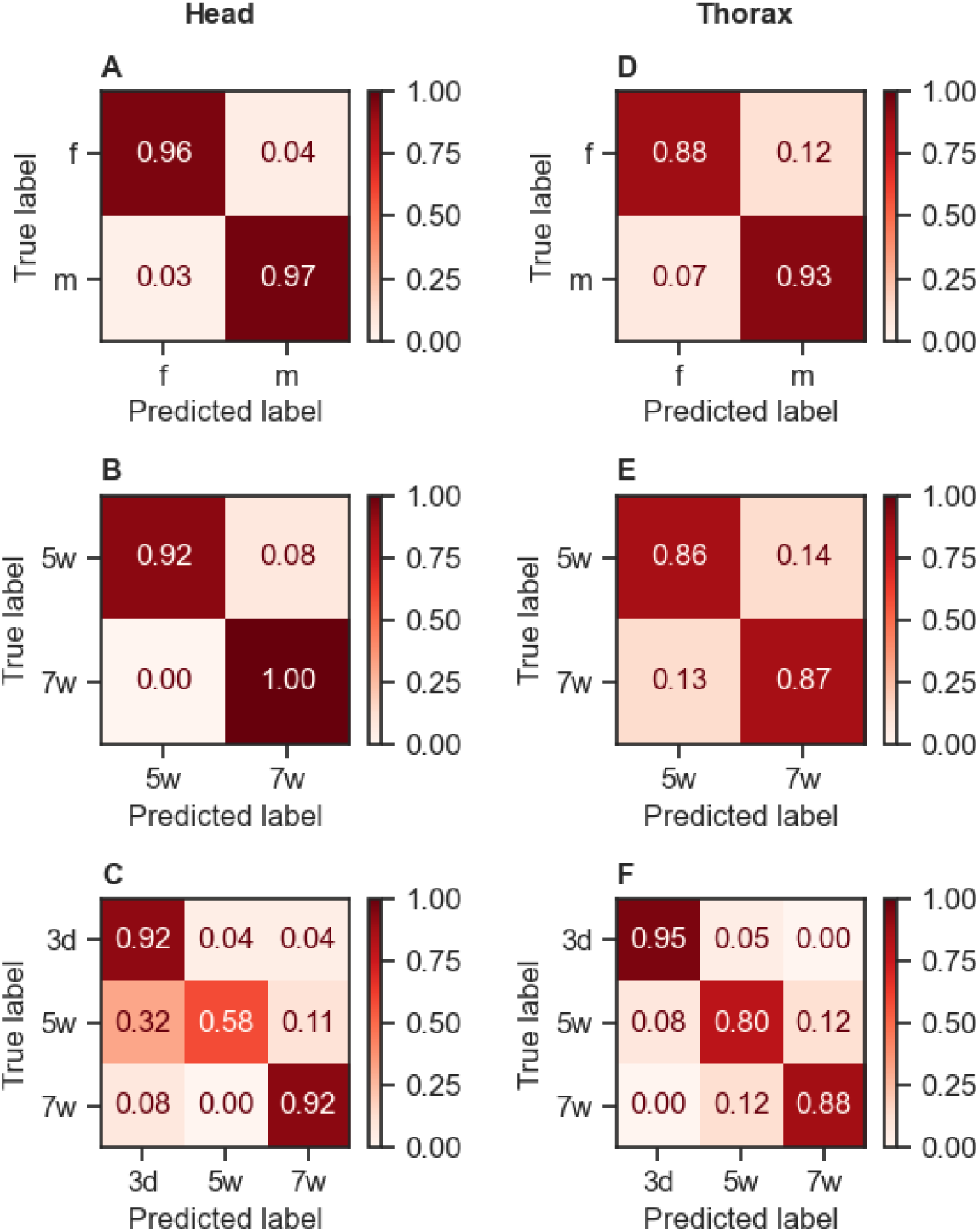
Confusion matrix for predicting tsetse sex and age using reduced number of wavenumbers. Accurate identification of females (f) and males (m) (**A, D**) and two-week age difference (5 weeks (5w) vs 7 weeks (7w) old) in male flies (**B, E**). Spectra from the thoraces of young female flies (3d post emergence) compared to older female flies (5 weeks (5w) and 7 weeks (7w) old (**C, F**)

**Fig 6.**
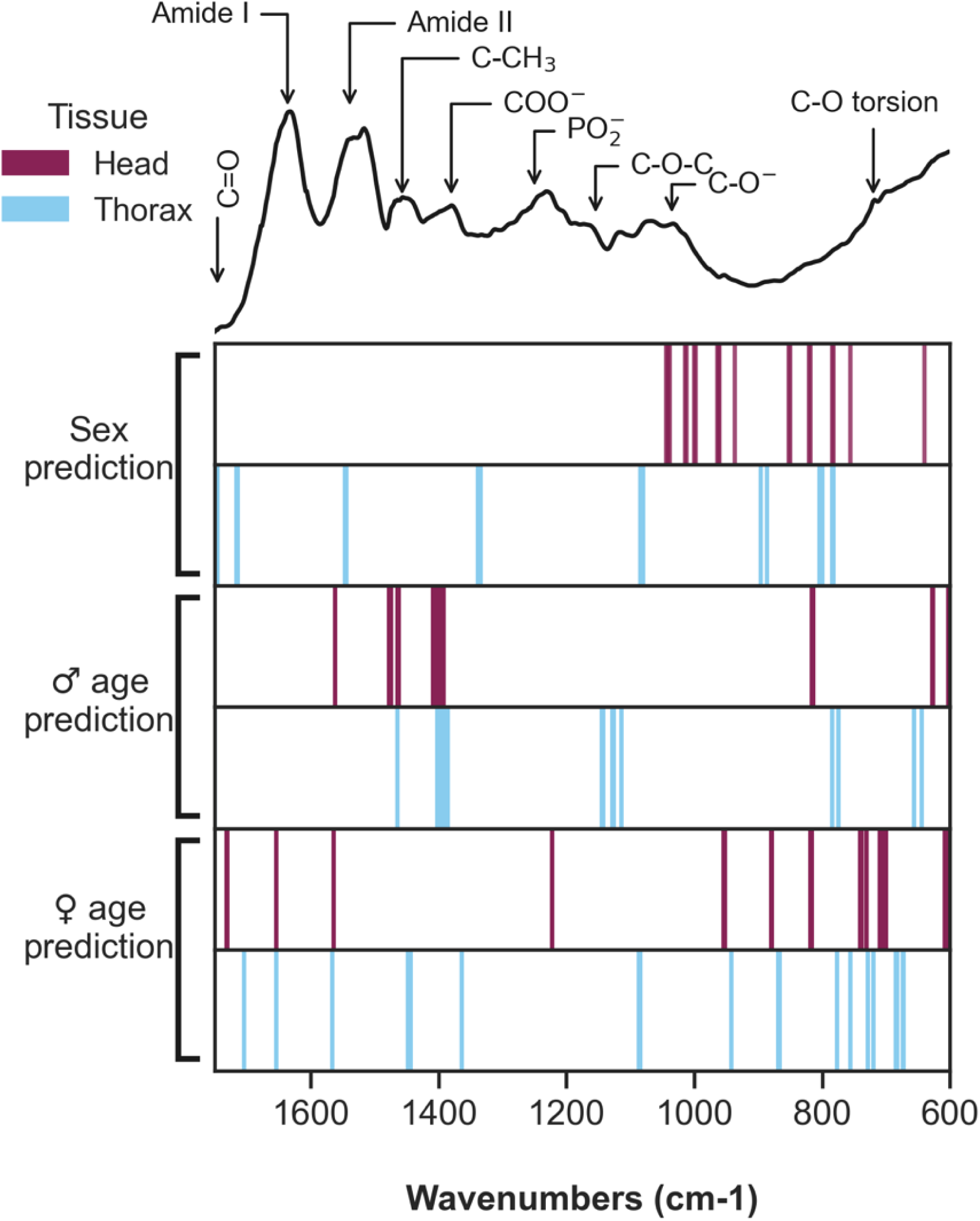
Important wavenumbers for predicting tsetse sex and age change depending on the trait predicted. Coloured lines represent the position of the most informative wavenumbers used by the models to predict sex, male age, and female age. Lines are coloured depending on the tissue used for MIRS: head (purple), thorax (light blue). Example spectra with band assignments is added on the top for reference.

**Table 3.** Accuracy, sensitivity and specificity of tsetse sex and age prediction on males and females in the training set and test set.

|  | Tissue | Accuracy<br>(train set) | Accuracy<br>(test set) |
| --- | --- | --- | --- |
| **Sex prediction** | Head | 0.95 ± 0.04 | 0.96 |
|  | Thorax | 0.93 ± 0.03 | 0.90 |
| **Males age prediction** | Head | 0.85 ± 0.06 | 0.94 |
|  | Thorax | 0.82 ± 0.06 | 0.86 |
| **Females age prediction** | Head | 0.84 ± 0.05 | 0.83 |
|  | Thorax | 0.85 ± 0.04 | 0.87 |

## Discussion

Here, we showed for the first time how a MIRS + ML toolbox can be applied to predict the sex and age of desiccated insectary-reared tsetse. The spectra collected from the head and thorax, but not the abdomen, allow accurate sex prediction. Age grading was successful in both sexes, even when flies were only two weeks apart in age. When using exclusively the thorax, this toolbox can easily differentiate between females and males using the infrared region related to lipids and carbohydrates. Interestingly, predictions using the head identified a narrow spectral region related to lipids as the most informative. It has been previously reported that *G. pallidipes* females possess a higher amount of cuticular lipids than males[22], which is likely linked to the female sex pheromone that constitutes the main cuticular hydrocarbon. Considering this potential bias, we did not mix the two sexes for analysis since the signal difference can mask the differences between the ages of each sex.

When analysing the most important regions for age grading in both males and females, some clear patterns emerged depending on the tissue and biological trait. In male flies, the C-CH_3_ and COO^-^ bands were consistently important in age grading for all tissues. However, the bands related to proteins and lipids and the –(CH_2_)-rock functional group related to wax was important across female tissues. Characterizing the informative and predominant wavenumbers is an important for understanding the association between age and absorption bands, which can be used to optimize data collection or model generalisation. An early staining method showed a relationship between cuticular layers in the thorax from laboratory and field caught flies[23]. Other methods using gene expression panels have also found that genes related to cuticular proteins were important for age grading. One study used RNAseq to analyse gene expression associated with age and sex in *G. m. morsitans* that were sourced from the same colony at LSTM [7]. Out of the ten genes shortlisted in the study, two proved to be enough for accurate age classification, one of these being cuticular protein 92F (GMOY002920). A second cuticular protein, 49Aa (GMOY005321), was also part of the list [10]. Previous work using MIRS with other insect vectors also reported differences in female cuticles between very young and old individuals, and the model predicted 3-day old females with minimal misclassification. However, when differentiating between 5- and 7-week-olds, the misclassification between both classes increased.

When we used the complete spectra for training, we found that LR and SVM with a linear kernel used the region from 2500 – 1800 cm^-1^ to predict sex and age, which does not contain any biochemical information related to insect cuticle. To ensure the algorithms learn from the biochemical differences between sexes and age groups, we restricted the inputs to specific spectral regions and limit the features the model uses. The strength of machine learning lies in finding patterns to separate classes; however, patterns can arise from confounding effects of contamination by water and CO_2_ rather than from the structural constituents of the specimen. It is important to diagnose and assess what the model is learning to rule out any bias and avoid overfitting. In spectroscopy data, variation between samples (i.e., baseline offset, variation on CO_2_ levels during different days when measuring) was robust enough for the model to accurately classify age and sex.

When determining the feasibility of using different tsetse tissues for analysis, the abdomen showed inconsistent spectra compared to the head and thorax, which might be caused by the presence of blood from previous meals and incomplete desiccation. However, the information from tsetse abdomens could still be used to identify blood meal sources, as demonstrated by the application of MIRS with *Anopheles* mosquitoes [24]

In summary, our results provide proof-of-principle for how MIRS can detect cuticular signals linked to ageing in tsetse. Future validation of this technique using field samples is needed, where environmental cues (naturally minimised in housed insect colonies) impact ageing rates. The next step will be to test the MIRS toolbox against wild tsetse collected from endemic areas, and preferably a region currently implementing vector control strategies. The machine learning models we describe here need to be further refined using more insectary-reared flies alongside a small complementary set of field samples (age-graded when trapped) to be able to confirm the efficacy and accuracy of this technology in the field [12].

## Conclusions

Our data strongly support the use of MIRS for high-accuracy age grading of both male and female *Glossina spp.* reared under insectary conditions. The protocol’s robustness, minimal maintenance, cost-effectiveness, and speed make it an ideal technique for vector surveillance programmes in resource-limited settings, and implementation will strengthen ongoing control efforts to control transmission of African trypanosomiasis.

## Acknowledgements

We are grateful to Jonathan Thornton, the dedicated technician overseeing the tsetse colony at LSTM, for his invaluable assistance in procuring tsetse flies for this work.

## Author contributions

Conceptualization: F.B., L.R.H.

Data curation: M.P., K.M.S.

Formal analysis: M.P

Funding acquisition: F.B, L.R.H.

Investigation: M.P, I.C, K.M.S.

Methodology: F.B., L.R.H.

Project administration: F.B., L.R.H.

Resources: F.B., L.R.H.

Supervision: F.B., L.R.H

Visualization: M.P, K.M.S.

Writing – original draft: M.P.

Writing – review & editing: M.P, K.M.S., I.C, S.A.B., F.B, L.R.H.

