## Supplementary information for "Advancing age grading techniques for *Glossina morsitans morsitans*, vectors of African trypanosomiasis, through mid-infrared spectroscopy and machine learning"

**Table S1**. Number of tsetse used for training the algorithm and creating test sets.

| Classification problem | Tissue | Dataset | # Samples | class/# samples | |
|---|---|---|---|---|---|
| **Sex prediction** | Head | Training | 425 | ♂ | 156 |
| | | | | ♀ | 269 |
| | | Test | 107 | ♂ | 31 |
| | | | | ♀ | 76 |
| | Thorax | Training | 431 | ♂ | 150 |
| | | | | ♀ | 281 |
| | | Test | 108 | ♂ | 35 |
| | | | | ♀ | 73 |
| **Age prediction females** | Head | Training | 276 | 3 days | 109 |
| | | | | 5 weeks | 73 |
| | | | | 7 weeks | 73 |
| | | Test | 69 | 3 days | 24 |
| | | | | 5 weeks | 19 |
| | | | | 7 weeks | 26 |
| | Thorax | Training | 283 | 3 days | 114 |
| | | | | 5 weeks | 71 |
| | | | | 7 weeks | 98 |
| | | Test | 71 | 3 days | 22 |
| | | | | 5 weeks | 25 |
| | | | | 7 weeks | 24 |
| **Age prediction males** | Head | Training | 149 | 5 weeks | 68 |
| | | | | 7 weeks | 81 |
| | | Test | 38 | 5 weeks | 26 |
| | | | | 7 weeks | 12 |
| | Thorax | Training | 148 | 5 weeks | 71 |
| | | | | 7 weeks | 71 |
| | | Test | 37 | 5 weeks | 22 |
| | | | | 7 weeks | 15 |

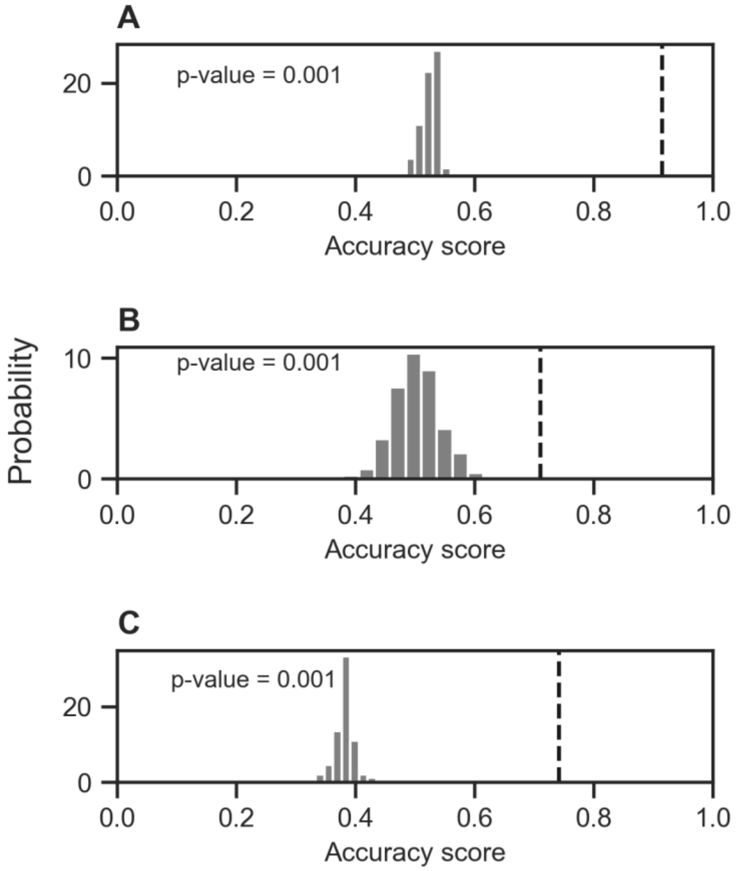


**Fig S1. Real dependency exists between features and different classes**. Labels were permutated while the features remained the same. Then, a model was fitted, and the accuracy was calculated. This process was repeated 1000 times. Histograms show the distribution of those accuracy scores using MIRS measurements from heads. Black dashed line indicates the accuracy score using the data without label permutation. P-values indicate permutated and non-permutated accuracies were significantly different. Accuracy scores for **A**. Sex prediction, **B.** Age prediction using males and **C.** Age prediction using females.

**Table S2.** Accuracy, sensitivity and specificity of tsetse sex and age prediction on males and females in the training set and test set using the whole spectral data

| | Tissue | Accuracy (train set) | Accuracy (test set) |
|---|---|---|---|
| **Sex prediction** | Head | 0.94 ± 0.02 | 0.99 |
| | Thorax | 0.94 ± 0.03 | 0.94 |
| **Males age prediction** | Head | 0.85 ± 0.06 | 0.89 |
| | Thorax | 0.88 ± 0.05 | 0.92 |
| **Females age prediction** | Head | 0.86 ± 0.04 | 0.93 |
| | Thorax | 0.92 ± 0.05 | 0.93 |

**Table S3.** Baseline performance of three machine learning models with different algorithms and pattern analysis (kernels) using different regions of the infrared range. The models are Support vector machine (SVM) with two kernels: radial basis function (rbf) and linear, Logistic Regression (LR) and Random Forest (RF). Each performance was measured using different regions of the mid-infrared spectra: 4000 – 600 $cm^{-1}$ (D1), 1800 – 600 $cm^{-1}$ (D2), 3500 – 2500 $cm^{-1}$ (D3), 2500 – 1800 $cm^{-1}$ (D4), 3500 – 2500 $cm^{-1}$ and 1800 – 600 $cm^{-1}$ combined (D5).

| | | | | 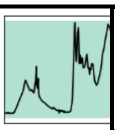 | 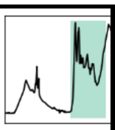 | 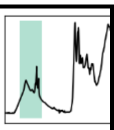 | 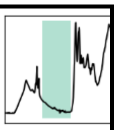 | 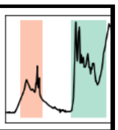 |
|---|---|---|---|---|---|---|---|---|
| | | **Model** | **kernel** | **D1** | **D2** | **D3** | **D4** | **D5** |
| **Male age prediction** | Thorax | SVM | rbf | 0.72 | 0.81 | 0.60 | 0.56 | 0.78 |
| | | SVM | linear | 0.94 | 0.90 | 0.85 | 0.94 | 0.88 |
| | | LR | -- | 0.89 | 0.86 | 0.80 | 0.91 | 0.84 |
| | | RF | -- | 0.72 | 0.76 | 0.65 | 0.59 | 0.76 |
| | Head | SVM | rbf | 0.71 | 0.66 | 0.67 | 0.66 | 0.68 |
| | | SVM | linear | 0.89 | 0.86 | 0.81 | 0.88 | 0.87 |
| | | LR | -- | 0.90 | 0.84 | 0.81 | 0.83 | 0.84 |
| | | RF | -- | 0.69 | 0.73 | 0.68 | 0.64 | 0.68 |
| **Female age prediction** | Thorax | SVM | rbf | 0.75 | 0.77 | 0.72 | 0.63 | 0.78 |
| | | SVM | linear | 0.90 | 0.86 | 0.86 | 0.89 | 0.85 |
| | | LR | -- | 0.91 | 0.87 | 0.85 | 0.89 | 0.88 |
| | | RF | -- | 0.77 | 0.77 | 0.71 | 0.67 | 0.78 |
| | Head | SVM | rbf | 0.80 | 0.80 | 0.74 | 0.72 | 0.80 |
| | | SVM | linear | 0.89 | 0.85 | 0.85 | 0.85 | 0.85 |
| | | LR | -- | 0.89 | 0.86 | 0.83 | 0.87 | 0.87 |
| | | RF | -- | 0.78 | 0.82 | 0.79 | 0.71 | 0.82 |
| **Sex prediction** | Thorax | SVM | rbf | 0.92 | 0.93 | 0.91 | 0.92 | 0.91 |
| | | SVM | linear | 0.97 | 0.95 | 0.95 | 0.95 | 0.97 |
| | | LR | -- | 0.95 | 0.95 | 0.94 | 0.95 | 0.96 |
| | | RF | -- | 0.91 | 0.92 | 0.89 | 0.91 | 0.90 |
| | Head | SVM | rbf | 0.89 | 0.88 | 0.91 | 0.87 | 0.90 |
| | | SVM | linear | 0.96 | 0.94 | 0.94 | 0.95 | 0.96 |
| | | LR | -- | 0.95 | 0.94 | 0.94 | 0.94 | 0.95 |
| | | RF | -- | 0.90 | 0.89 | 0.91 | 0.89 | 0.92 |

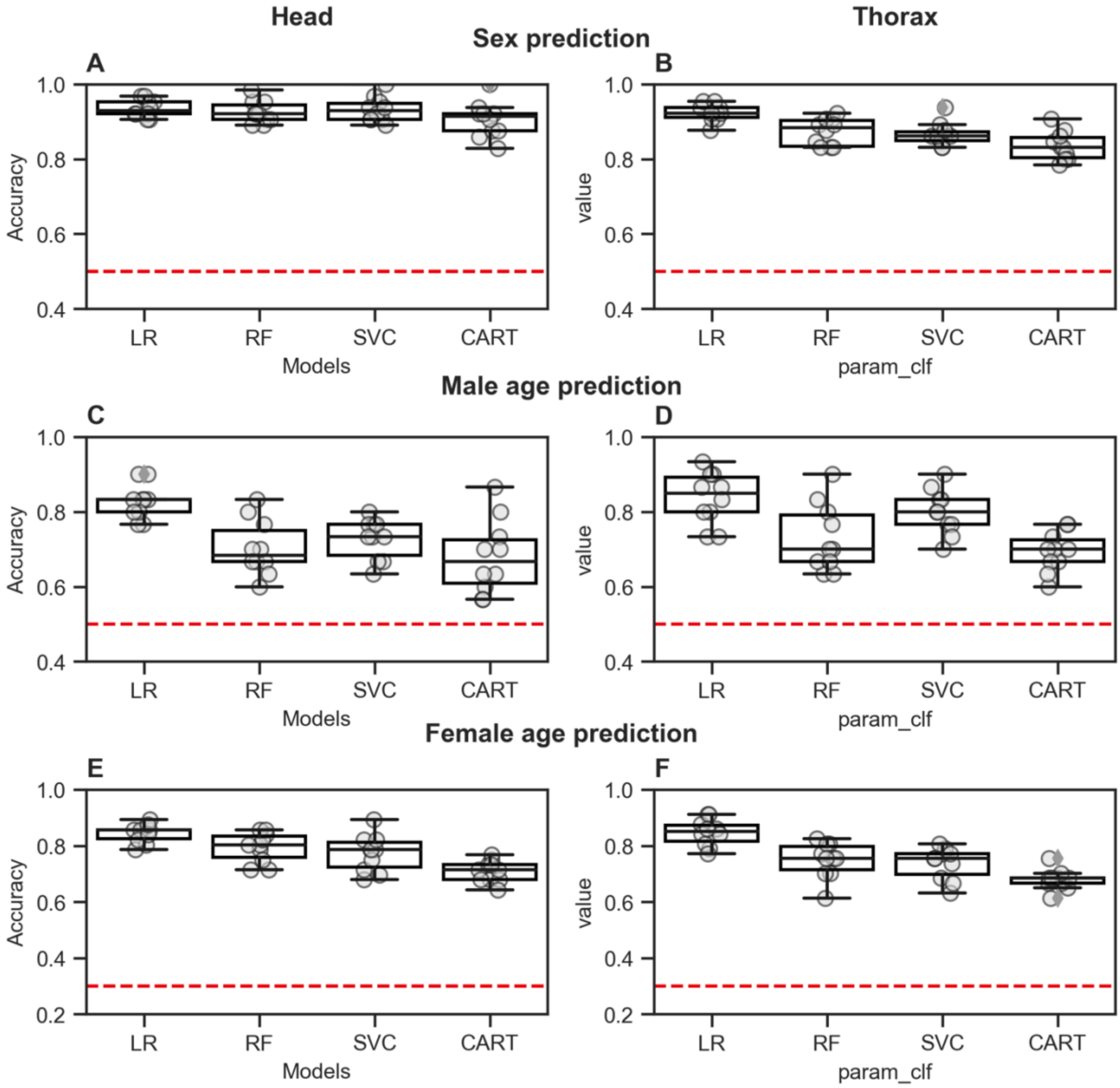


**Fig S2. Model performance is maintained with reduced wavenumbers.** Model performance on training set of various ML models (LR: Logistic regression, RF: random forest, SVC: support vector machine and CART: decision tree classifier) using the region between 1800 to 600 $cm^{-1}$. Spectral data from the heads (left column) and thorax (right column) for identification of sex (**A, B**), age of males (**C, D**) and females (**E, F**). Boxplots show the distribution of accuracies after 10-fold cross-validation. Red line shows the accuracy of a random model (0. (0.5 for binary classifications, 0.3 for three-way classifications) Each dot represents the accuracy of one-fold.

**Table S4.** Most important wavenumber regions for sex/age prediction and their assignment

| Biological trait | Tissue | Wavenumbers ($cm^{-1}$) | Functional group assigned(1–4) | Reference |
|---|---|---|---|---|
| Sex identification | Head | 1044 1042 1040 1038 | C-O-P stretching | Lipids |
| | | 1014 1012 1000 998 964 962 960 | C-O stretch | Chitin, protein, wax |
| | | 936 | * | * |
| | | 852 850 | * | * |
| | | 820 818 | $PO_2^-$ symmetric stretching | Lipids |
| | | 784 782 756 | * | * |
| | | 640 | * | * |
| | Thorax | 1750 1748 1746 1716 1714 | C=O stretching | Lipids, waxes, proteins |
| | | 1546 1544 | Amide II | Proteins |
| | | 1338 1336 1334 | C-OO- | Proteins, chitin |
| | | 1084 1082 1080 | C-O stretch | Lipids |
| | | 896 886 804 802 800 784 782 | * | * |
| Males age identification | Head | 1562 | Amide II | Proteins |
| | | 1478 1476 1474 1464 1462 | C-H bend | Lipids |
| | | 1408 1406 1402 1400 1398 1396 1394 1392 | C-H bend | Proteins |
| | | 816 814 628 626 | * | * |
| | | 602 | C=O bending | Proteins |
| | Thorax | 1750 | C=O | Lipids, wax |
| | | 1464 | C-H | Lipids, wax |
| | | 1402 1400, 1398 1396 1394 1392 1390 1388 1386 | C-H | Proteins |
| | | 1144 1142 1128 1126 1114 | C-O stretch | Chitin |
| | | 784 774 | * | * |
| | | 656 644 | * | * |
| Females age identification | Head | 1732 1730 | C=O stretch | Proteins, waxes |
| | | 1654 | Amide I | Proteins, chitin |
| | | 1564 | Amide II | Proteins, chitin |

| | | | | |
|---|---|---|---|---|
| | | 1222 | C-H | Proteins, wax |
| | | 954 952 | C-O stretch | |
| | | 880 878 818 816 | * | * |
| | | 740 738 730 | -CH2-rock | |
| | | 710 708 702 700 | C-O torsion | |
| | | 608 606 | C=O bending | Proteins |
| | | 1704 | C=O | Lipids, wax |
| | | 1654 | Amide I | Proteins, chitin |
| | | 1566 | Amide II | Proteins, chitin |
| | | 1448,1446,1444, | CH2 bending | Lipids, proteins |
| | | 1364 | C-H | Wax, proteins |
| | **Thorax** | 1086,1084 | $PO_2^-$ symmetric stretching | Lipids, nucleic acids |
| | | 942 | * | * |
| | | 868,866, 776,756,728 | * | |
| | | 720, 684,682, | C-O torsion | * |
| | | 674,672 | * | * |

*No known assignment
